# scTrimClust: A Fast Approach to Robust scRNA-seq Analysis Using Trimmed Cell Clusters

**DOI:** 10.1101/2025.04.16.649082

**Authors:** Sergej Ruff, Klaus Jung

## Abstract

Detection of marker genes and other data analyses in single cell RNA-sequencing (scRNA-seq) experiments very much rely on the result of unsupervised clustering of cells. However, in 2-dimensional representation of clustering results, several cells appear as outliers or in the border area of a cluster suggesting that these cells may not adequately represent a particular cell type.

We propose a novel and fast approach, scTrimClust, for identifying cells that may be interpreted of extreme specimens of their cell type. Identification is based on concave hulls build around each 2-dimensional cell cluster and the distance of each cell to the border area of its population.

We study in two data examples, how cells with non-representative expression profile can influence the results of the analysis. We found that some sets of marker genes are little influenced by extreme cells while other sets are strongly modified, and must therefore be treated carefully.

scTrimClust is also useful to compare the influence of other parameters of an scRNA-seq analysis, e.g. normalization or the clustering approach, on the results. We also provide a software implementation of scTrimClust.

## 1. Introduction

Single cell RNA-sequencing (scRNA-seq) (Kolodziejczyk et al., 2015) has become an important method in systems biology to analyse cell-type specific gene expression and has been applied to research on a wide range of diseases. For example, Gonzáles et al. (2020) used scRNA-seq to unravel tumor heterogeneity, and Melms et al. (2021) described the cell-type specific transcriptome in the lung under COVID-19 infections.

After mapping and counting of sequencing reads per gene, a high-dimensional expression matrix X with *d* rows representing several thousands of genes and *n* columns representing several hundred or thousands of individual cells from the tissue sample is the starting point of further analysis. Still, the number of rows exceeds usually the number of columns.

One fundamental step in the data analysis of scRNA-seq experiments is clustering and identification of cell types which can be done by unsupervised clustering methods such as the *k*-nearest neighbour (KNN) method based on the dimension-reduced expression matrix X. Depending on the type of tissue that is under research, clustering results in several more frequent cell types with several hundred cells and several less frequent cell types with one hundred cells of less.

While clustering itself is mostly done in the space of a few principal components, the clustering results are usually visualized in a 2-dimensional *t*-SNE (Van der Maaten and Hinton, 2008) of UMAP (McInnes et al., 2018) plot. The shape of the different cell clusters in these plots can be quite different with some clusters having a concave or extremely skewed shape and sometimes having clear outliers. This suggests that the membership of outlying cells or cells in the border area of a cluster might not perfectly represent the cell type. Although these cells can be interpreted simply as biological extremes which should be kept in the analysis, there is also a risk that these cells might be classified to the wrong cluster. Anyhow, these cells can have a strong impact on the results of the whole analysis.

To downweigh the effect of extreme observations, trimmed estimators have been used in statistics since the 1980s (Huber and Ronchetti, 1981). For example, the trimmed mean has been shown to be a robust estimator in the presence of extreme observations. In univariate or bivariate data, outliers or extreme observations can be identified by statistical tests or graphically, e.g. using box-and-whiskers-plots or bagplots (Kruppa and Jung, 2017). Following the idea of trimmed estimators, we introduce here a novel approach for fast identification of outlying cells and cells in the border area of each cell cluster. Since cell cluster in a t-SNE or UMAP plot often show a skewed or concave shape, existing methods for outlier detection are not appropriate. Therefore, using the coordinates of the *t*-SNE or UMAP plot together with the clustering results, the concave hull of each cluster is determined as a basis to determine the most representative cells per cluster.

The effect of extreme and near border cells can then be studied during the analysis. Taking the fact, that the results of scRNA-seq data analysis is influenced by several parameter settings, our approach is very helpful to study the robustness of the whole analysis. For example, the choice of the normalization method, of the number of principal components used during unsupervised clustering, and of gene filtering can impact the overall results. While robustness of a scRNA-seq analysis can also be studied using time-consuming bootstrap procedures, the novel approach presented here simply contrasts the whole data set versus a reasonably trimmed data set and is therefore much faster.

In this manuscript, we describe our novel approach for trimming sets of cell types based on the identification of extreme and near border cells. We demonstrate the practicability of our approach on two data examples: first the data set of Peripheral Blood Mononuclear Cells (PBMC) that is also used in the tutorial of the Seurat R-package (Satija et al., 2015), second a COVID-19 data set. We also provide a description of how to use our novel R-function that implement the scTrimClust approach.

## 2. Methods

### 2.1. Data examples: peripheral blood cells and whole blood samples

We evaluated our approach using two publicly available scRNA-seq data sets. The first one, names PBMC 3k, was generated from peripheral blood mononuclear cells, is featured in the Seurat guided tutorial and is accessible through the Seurat package or the 10x Genomics webpage (https://cf.10xgenomics.com/samples/cell/pbmc3k/pbmc3k_filtered_gene_bc_matrices.tar.gz). It includes 2,700 PBMCs with 13,714 features from healthy human donors.

The second data set, focusing on SARS-CoV-2 infection, contains whole blood samples depleted of red blood cells from 7 COVID-19-positive patients (stratified into mild, moderate, and severe groups) and from 3 SARS-CoV-2-negative controls. For our analysis, we retained only the 5,295 cells and 38,949 features from the 7 positive cases. This data set, originally generated by Silvin et al. (2020), is accessible via the ArrayExpress Archive of Functional Genomics Data under accession number E-MTAB-9221 (https://www.ebi.ac.uk/biostudies/arrayexpress/studies/E-MTAB-9221).

### 2.2. Standard workflow of scRNA-seq analysis

Both data sets are first analysed using a standard workflow with no cells excluded from the analysis, following the analysis pipeline of the Seurat R-package (Satija et al., 2015). This pipeline consists of first filtering cells with extremely low (due to cell damage) or extremely high (due to non-dissolved cells) feature counts followed by normalization. Thus, there is already some filtering regarding inappropriate data.

Thereafter, the data set is further filtered by excluding genes with low variability over all cells as these genes may have little biological meaning. Typically, less than 5,000 genes remain in the analysis. Next, linear dimension reduction by principal component analysis (PCA) is run and the unsupervised clustering is then performed on a couple of principal components. Results of the clustering is shown in *t*-SNE or UMAP plots. As final steps, marker genes per cluster are identified – usually the most strongly expressed genes per cluster – and differential expression analysis between the clusters in performed.

Using the scTrimClust approach, we compare the results of marker gene detection and of differential expression analysis under several different settings, i.e. with two normalization methods (LogNormalize versus CLR), with two different sets of preselected genes (1,000 versus 2,000), and with two different numbers of principal components (5 versus 10) used for the clustering step.

### 2.3. scRobust workflow of scRNA-seq analysis

To identify extreme cells per cluster, we determine the concave hull of each cluster in the 2-dimensional *t*-SNE coordinates obtained in the standard analysis. For that purpose, we use the function `ahull` of the R-package alphahull. This functions determines *T* cells of each cluster *k* as anker points 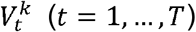 to span the concave hull. Next, we calculate the Euclidean distances 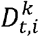 of each cell *i* in a cluster *k* to the anker points of the concave hull in the 2-dimensional *t*-SNE space. The minimum distance of each cell to one of the anker points, 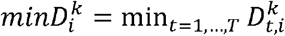, is used to judge whether this cell belongs more set of representative cells of to those in the border area. Results, e.g. sets of selected markers, can then be compared by excluding _α_% of the most extreme cells, i.e. those cells with 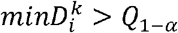, where Q_1-α_ is the (1 – _α_) – quantile of all *minD^k^* in cluster *k*.

### 2.4. Evaluation by set comparisons and breakdown point analysis

An essential goal of scRNA-seq is the selection of marker genes for individual cell types. Therefore, we compare the sets of selected markers under the different settings. For both data sets, COVID-19 and PMBP, we analysed how cells from the border areas influence the selection of marker genes under a) different normalization approaches, b) under different numbers of preselected genes, and c) under different numbers of PCs used for unsupervised clustering.

Under each scenario and for each cell type we determined the set S_full_ of marker genes obtained with the full data and the set S_trim_ of marker genes obtained with the trimmed data. From these two sets, we derived the sets

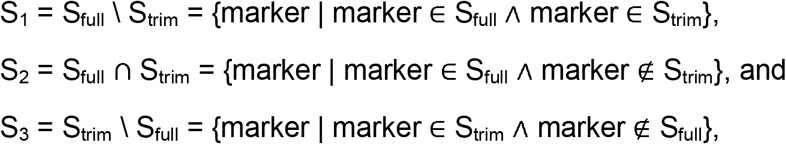

i.e., the differences and the intersect between S_full_ and S_trim_. Based on the cardinalities |S_1_|, |S_2_|, and |S_3_|, i.e. the number of markers in each set S_1_, S_2_ and S_3_, we calculated the percent proportions

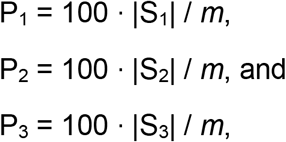

where *m* is the number of markers in the union of S_full_ and S_trim_, i.e., *m* = |S_full_ U S_trim_|.

In the case of large proportion P_2_, a finding can be seen as robust since full and trimmed data yield a highly similar set of markers. If, however, P_1_ is large, it means that the selected markers with the full data set highly rely on additional information from the cells in the border area. Finally, if P_3_ is large, it means that removing cells from the border area completely changes that set of selected markers.

In the area of robust statistics, estimators are often evaluated by their breakdown point, which is the proportion of extreme observations an estimator can cope with until it gives inaccurate results. Such a breakdown point analysis can also be done by comparing proportion P_1_ against P_2_ for different percentages of trimming. Hereby, the analyst can observe how robust a marker set is against the cells in the border area.

## 3. Results

### 3.1. Influence of cells from the border area on selection of marker genes

Here, we highlight exemplarily some scenarios with clear effects, further scenarios are presented as supplementary material.

In the PBMB data set, when the LogNorm normalization and 10 PCs where used, we compared the sets of selected markers when using the full data sets and the data when 10% of near border cells where removed. Figure 1 shows the *t*-SNE plot with concave hulls around each cluster. Furthermore, a heatmap shows the percentages P_1_, P_2_ and P_3_ for each cell type, separately for the scenarios with 1,000 and 2,000 preselected genes. For the largest clusters, representing Monocytes, T-cells and B-cells, P_2_ is close to 100% meaning that the marker sets S_1_ and S_3_ are highly similar and that the cells in the border area have little impact on the selection. In contrast, for NK-cells P2 is much smaller – around 60% – meaning a high uncertainty in the markers selected with the non-trimmed data. For low frequent cell types, no markers where selected after trimming (not shown in the heatmap).

**Figure 1:**
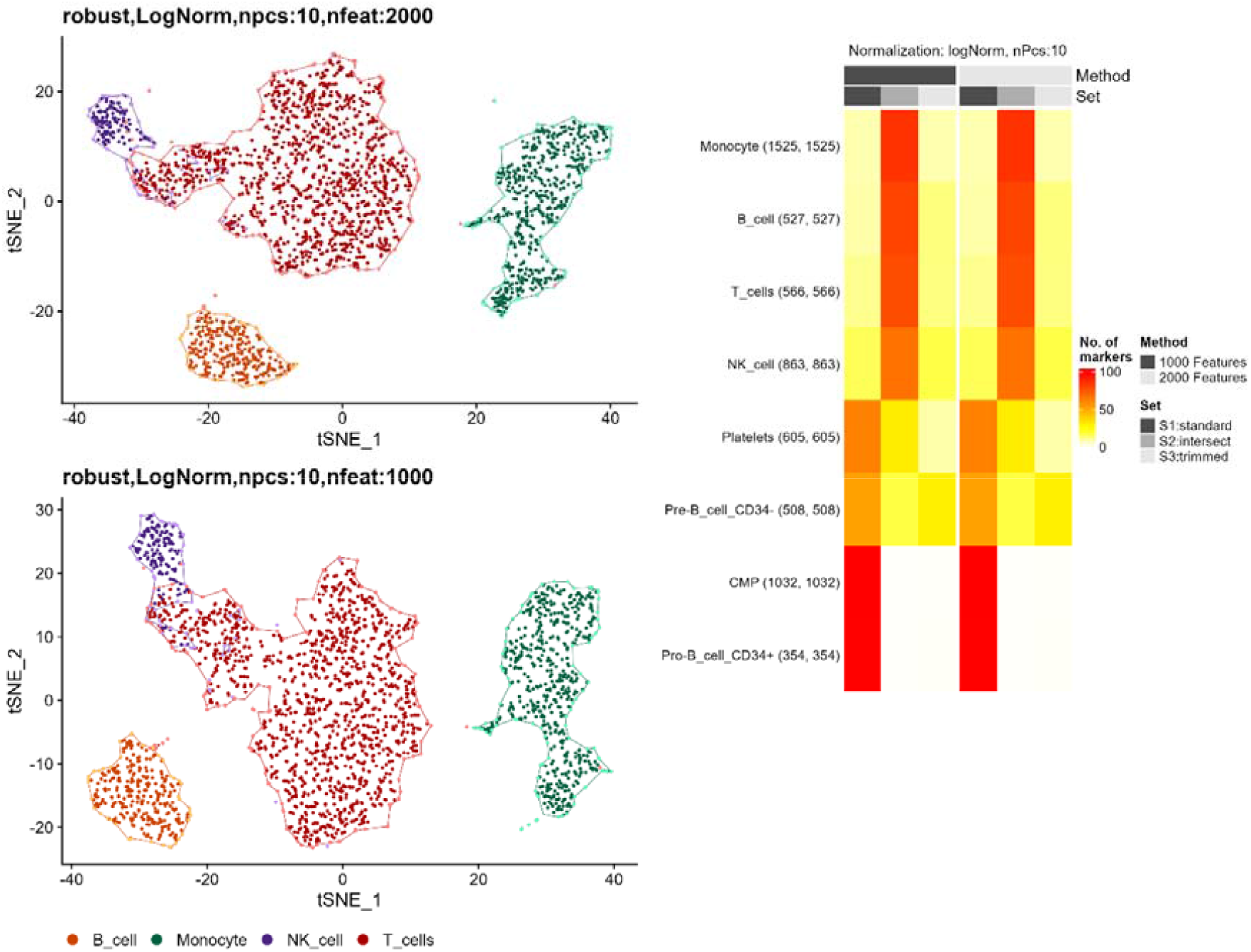
Left: *t*-SNE plot of the PBMB data set with convex hulls around each cell cluster. Right: Heatmap comparing results when using either 1,000 or 2,000 preselected genes (first three versus last three columns). While overlaps P_2_ are large for Monocytes, T-cells and B-cells, the overlap is much smaller for NK-cells.

The heatmap can also be used to compare the influence of using either 1,000 or 2,000 preselected features. Here, the patterns are mostly the same. Therefore, the number of preselected features appears to be not relevant.

In the COVID-19 data set, we examined the effect on robustness, when using 5 PCs and 2,000 preselected features at a higher trimming percentage of 40%. Here, T-cells exhibited the most substantial disparity in the number of detected marker genes, with 10,147 marker genes identified after CLR normalization compared to 6,417 after applying LogNorm (Figure 2). Furthermore, T-cells displayed the highest intersect percentage (P2)—92% under CLR versus 84% under LogNorm.

**Figure 2:**
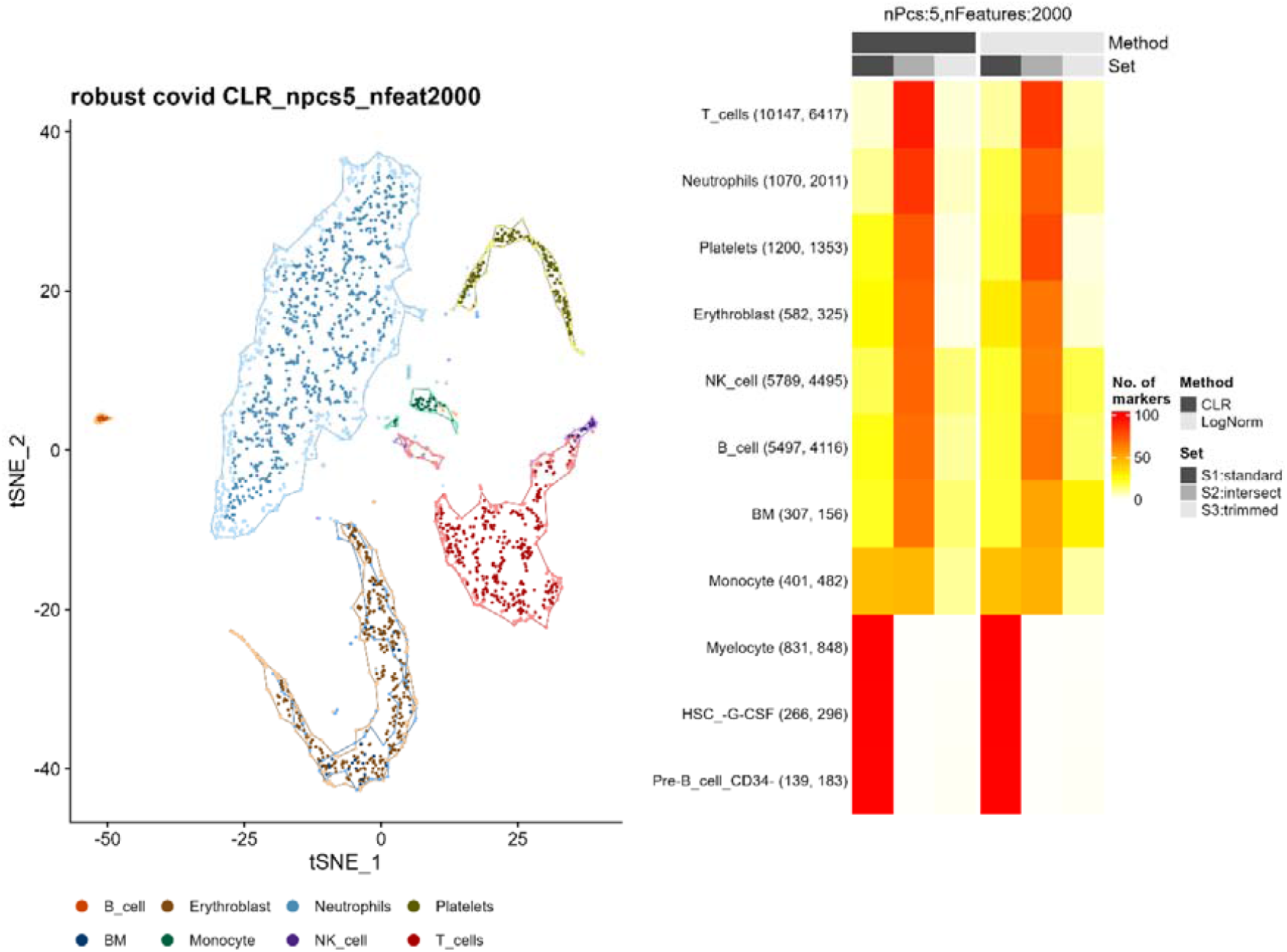
Left: *t*-SNE plot of the COVID data set with convex hulls around each cell cluster, where only the most frequent cell types are shown. Right: Heatmap comparing results under CLR and LogNorm normalization. The first three columns show high overlap between analyses with full and trimmed data sets under CLR normalization for T-cells, neutrophils, erythroblasts, NK cells, B-cells, and unspecific bone marrow cells. The last three columns show that more marker genes are retained for platelets and monocytes under LogNorm normalization, while the S2:intersect (P2) values for other cell types are slightly lower in comparison to CLR normalization. Cell types with the lowest number of cells retained no marker genes after trimming.

Beyond T-cells, CLR demonstrated consistently higher P2 percentages across other major cell types. For neutrophils, CLR achieved an intersect rate of 84%, exceeding LogNorm’s 72%. Similarly, erythroblasts showed a 71% overlap with CLR compared to 65% with LogNorm, while NK cells exhibited 70% (CLR) versus 62% (LogNorm). For B cells, CLR normalization maintained a slight advantage (68% versus 66%). Notably, unspecified bone marrow cells revealed a more pronounced difference, with 65% of marker genes conserved under CLR normalization relative to 51% under LogNorm normalization. LogNorm outperformed CLR in only two cell clusters: Platelets exhibited a marginally higher P2 value (78% for LogNorm vs. 75% for CLR), and monocytes showed a slight increase (48% vs. 46%). No marker genes were found for low frequent cell types.

### 3.2. Breakdown point analysis

As expected, marker selection is more stable for more frequent cell types. In both COVID and PBMC data sets—analyzed using CLR normalization, 5 principal components, and 1,000 preselected genes—low-frequency cell types failed to yield any detectable markers even at a minimal trimming level of 10% (Figure 3). Thus, 10% would be the breakdown point for these cell types.

**Figure 3:**
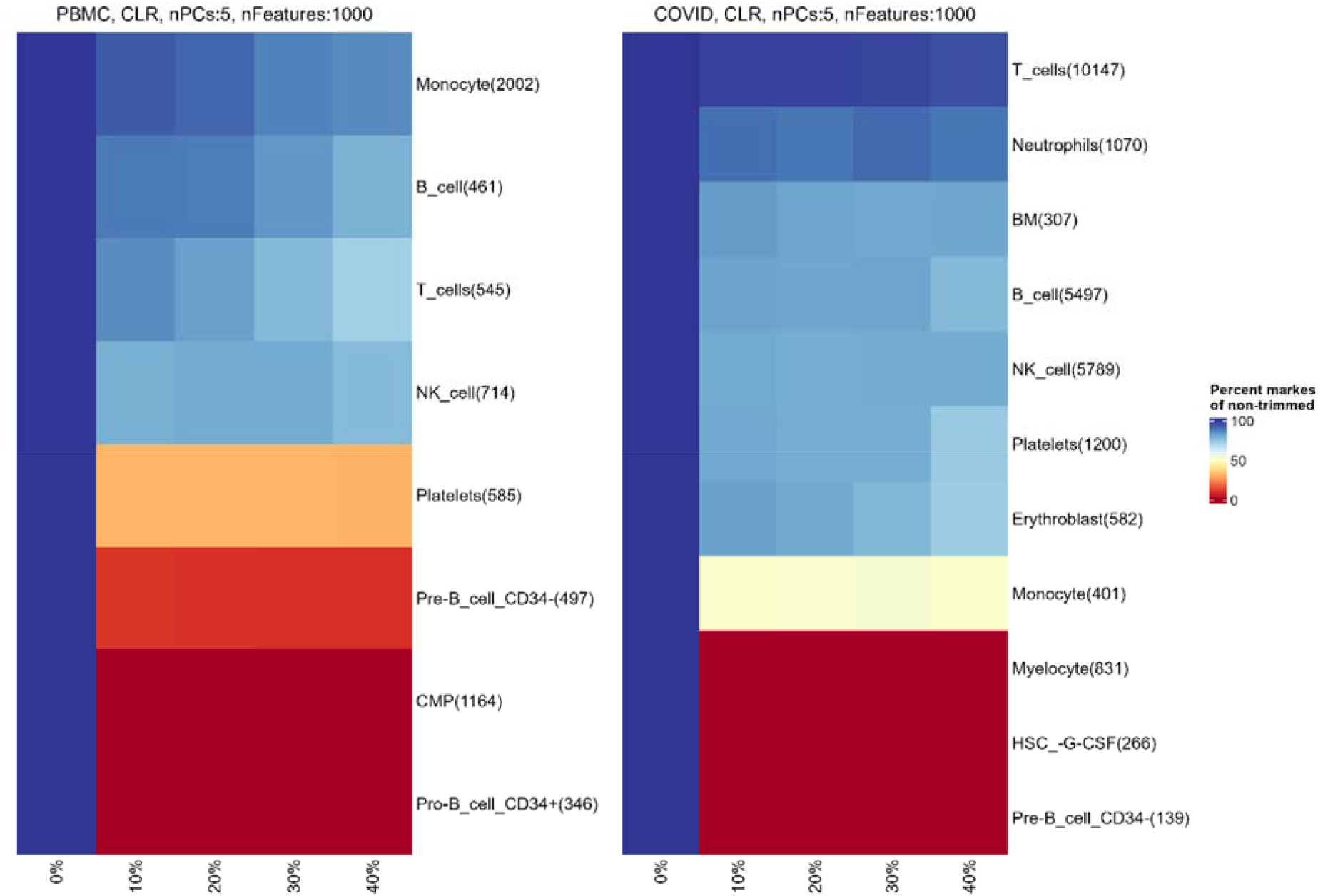
Left: Breakdown point heatmap of PBMCs (CLR-normalized, 5 PCs, 100 features). Right: COVID-19 data set under identical parameters, with color gradient showing percentage of retained marker genes (red:0% to blue:100%) after trimming at 0%, 10%, 20, 30%, and 40% thresholds, with y-axis labeling cell type clusters.

For more abundant cell types, the number of retained markers gradually decreased as trimming percentages increased. In the PBMC data set, marker gene retention varied across cell types with increasing thresholds. Monocytes retained 95% of their marker genes at 10% trimming, declining gradually to 92% (20%), 88% (30%), and 85% (40%). B cells, however, showed stable retention at 88% for both 10% and 20% trimming, followed by a drop to 83% (30%) and 78% (40%). This suggests that the initial shift from no trimming to 10% had the greatest impact, with retention stabilizing between 10% and 20% before further declines at higher trimming levels. The same plateau can be observed for the NK cell cluster. For platelets and CD34-negative pre-B cells, marker gene retention drops below 50% at just 10% trimming and remains stable at higher thresholds. Meaning, the breakdown point here is already hit at 10% trimming and further trimming does not change the number of marker genes significantly. A similar pattern is observed in the COVID-19 data set, where the monocyte cluster also stabilizes at 50% retention across trimming levels. The T-cell cluster maintains notably higher retention, exceeding 95% at every trimming threshold. For other abundant cell types with retention rates above 50%, values remain around 80%, with only minor decreases under more aggressive trimming.

## 4. Implementation in R

We added the approach of scTrimClust in form of three new functions to our R-package RepeatedHighDim (https://cran.r-project.org/web/packages/RepeatedHighDim/index.html). Here, we present these three functions with their main arguments. A more detailed description of how to use the functions is given in the supplementary material.

### 4.1. Identification of cells in the border area

The core function, `scTrimClust`, visualizes cell clusters in a low-dimensional space (t-SNE, UMAP, etc.) and removes cells from the border are of each cluster.

The `hull.alpha` parameter controls the concavity of cluster boundaries using a convex hull. Higher values produce smoother, more inclusive hulls, while lower values create tighter, more irregular contours. The `outlier.quantile` parameter sets the percentile cutoff (0-1) for minimum cell-to-hull distances, classifying cells below this threshold as extremes. Lower values restrict detection to extreme outliers, whereas higher values identify more peripheral cells as outliers.

`scTrimClust` returns a list of objects. The output includes a modified plot with flagged or removed outlier cells, along with the hull coordinates required to generate cluster boundaries, a list of ahull objects for each cluster, and the coordinates for both non-outlier and outlier cells. It also returns the Seurat object, with outlier cells removed if `remove.outliers=TRUE`. The processed Seurat object enables seamless integration of scTrimClust into existing Seurat workflows, where DimPlot would typically be used, allowing users to continue downstream analyses without outliers.

### 4.2. Comparative visualization of marker selection

The `scTC_trim_effect` function quantifies changes in cluster-specific marker genes after outlier removal by comparing untrimmed (default Seurat) and trimmed (scTrimClust-processed) data sets. It takes a list of marker genes produced by `Seurats FindAllMarkers` function. The output is a heatmap displaying cell clusters (rows) and three categories per method (columns): markers exclusive to untrimmed data, shared markers, and markers exclusive to trimmed data. An example is given in Figure 1 and 2. Colors represent the percentage of markers in each category (0-100%).

While `scTC_trim_effect` compares marker sets across methods at a fixed trimming level, `scTC_bpplot` evaluates marker retention across trimming percentages for a single method. Trimming percentages are specified via the `outlier.quantile` parameter in `scTrimClust`. The heatmap displays cell clusters (rows, labeled with original marker counts) and trimming percentages, using a color gradient (default: 0% = blue, 100% = red) to show the percentage of original markers retained. High retention (warmer colors) indicates robust clusters, while cooler colors reflect marker loss after trimming. An example is given in Figure 3.

## 5. Discussion

Advances in sequencing technology provide deeper insights into molecular mechanisms, however, as larger data sets arise from these experiments, more uncertainty is introduced as well. Uncertainty can come from the complex bioinformatics pipelines for omics analyses which consist of many steps where the analyst has to draw decisions about parameter settings in nearly each step. Here, we focus on the uncertainties in the data itself by trying to identify cell specimens which may not perfectly represent a particular cell type. This contrasts our approach with time-consuming resampling approaches which put each single observation on the scale. Though bootstrap and other resampling approaches have a long tradition to evaluate the robustness of findings in omics data (Pepe et al., 2003; Saremi et al., 2021; Hemandhar Kumar et al., 2024), they treat each observation to the same degree and can be very time-inefficient for large data sets and complex analysis pipelines. By focusing on the identification of extreme observations, we follow more the concept of trimmed statistics. To evaluate the robustness of a finding, the analyst must run the whole analysis only twice: once for the full data set and once for the trimmed one. Hubert et al. (2005) also used the idea of downweighing outliers in transcriptomics data when doing PCA.

We have demonstrated that our approach is also useful to compare the robustness between two variants of the analysis, e.g. two different normalization methods or two different approach in the clustering step. Sun et al. (2019) have also shown how sensitive scRNA-seq is to the choice of the clustering method. Therefore, by trimming cells with potential wrong cluster membership, we can study the effect of the extreme or outlying cells of a cluster.

As most computational methods, scTrimClust has some limitations, which we are currently working on to diminish. The detection of cells in the border area of a cluster itself depends on the clustering method and the methods for dimension reduction, thus on prior steps on the analysis. This, however, appears to be not a big issue, since the approach is still useful to compare the robustness of selected markers. In contrast, our approach is little useful for less frequent cell types. Trimming cells from a small population decreases statistical power and may lead to a higher false negative rate. Our approach also struggles a bit for cluster which appear with dislocated subclusters. The concave hull can than be split into disjunct parts and identification of near border cells does not perfectly work.

