## Supplementary Material for "scTrimClust: A Fast Approach to Robust scRNA-seq Analysis Using Trimmed Cell Clusters"

A) **Supplementary Figures**

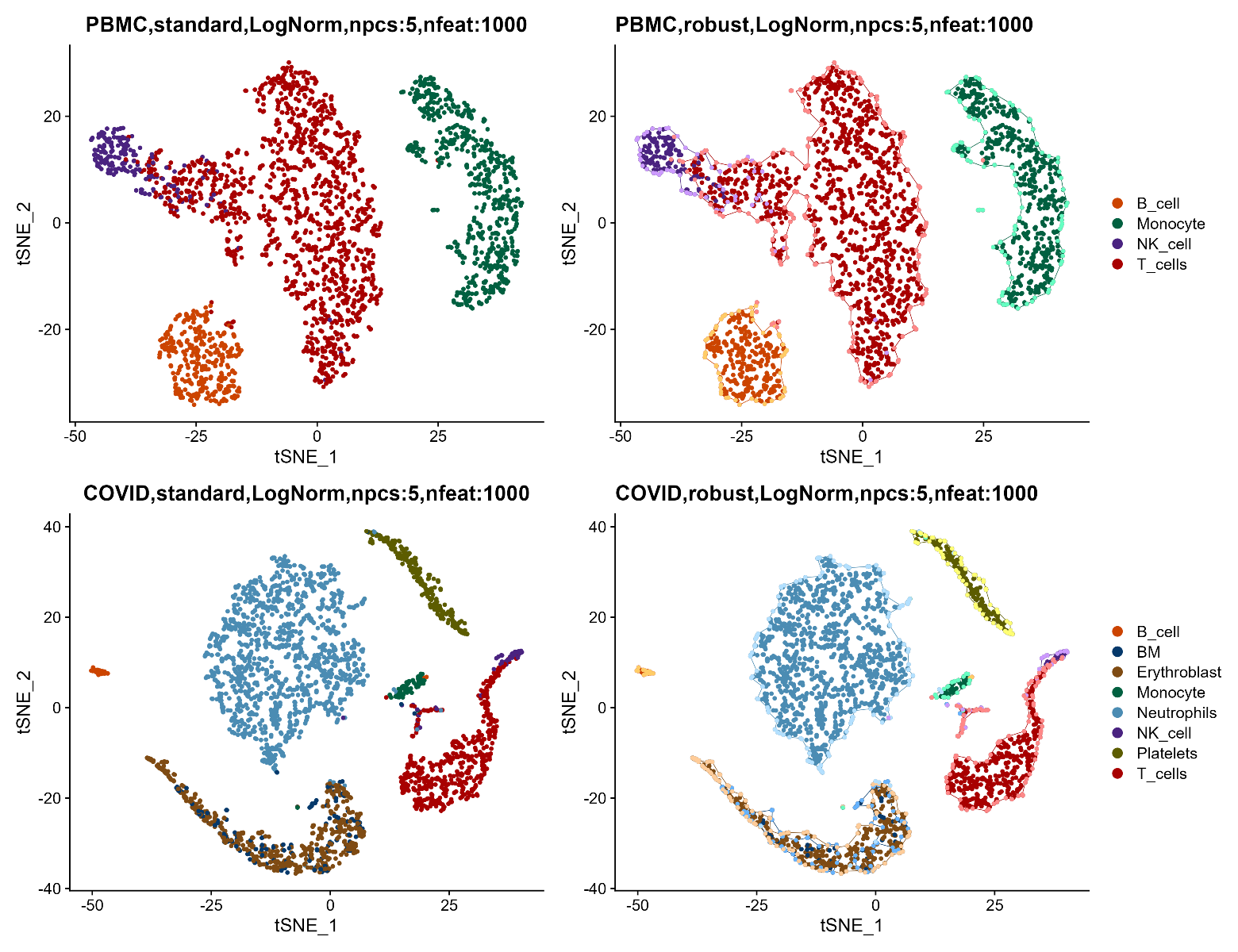

**Supplementary Figure 1:** t-SNE visualization of PBMC and COVID-19 datasets. Top-left: PBMCs plotted using Seurat’s DimPlot. Top-right: Same PBMC data processed with scTrimClust, displaying convex hulls around clusters and highlighting outliers (bright colors). Bottom-left: COVID-19 data with default DimPlot. Bottom-right: COVID-19 data analyzed with scTrimClust, showing enhanced cluster separation and outlier detection.

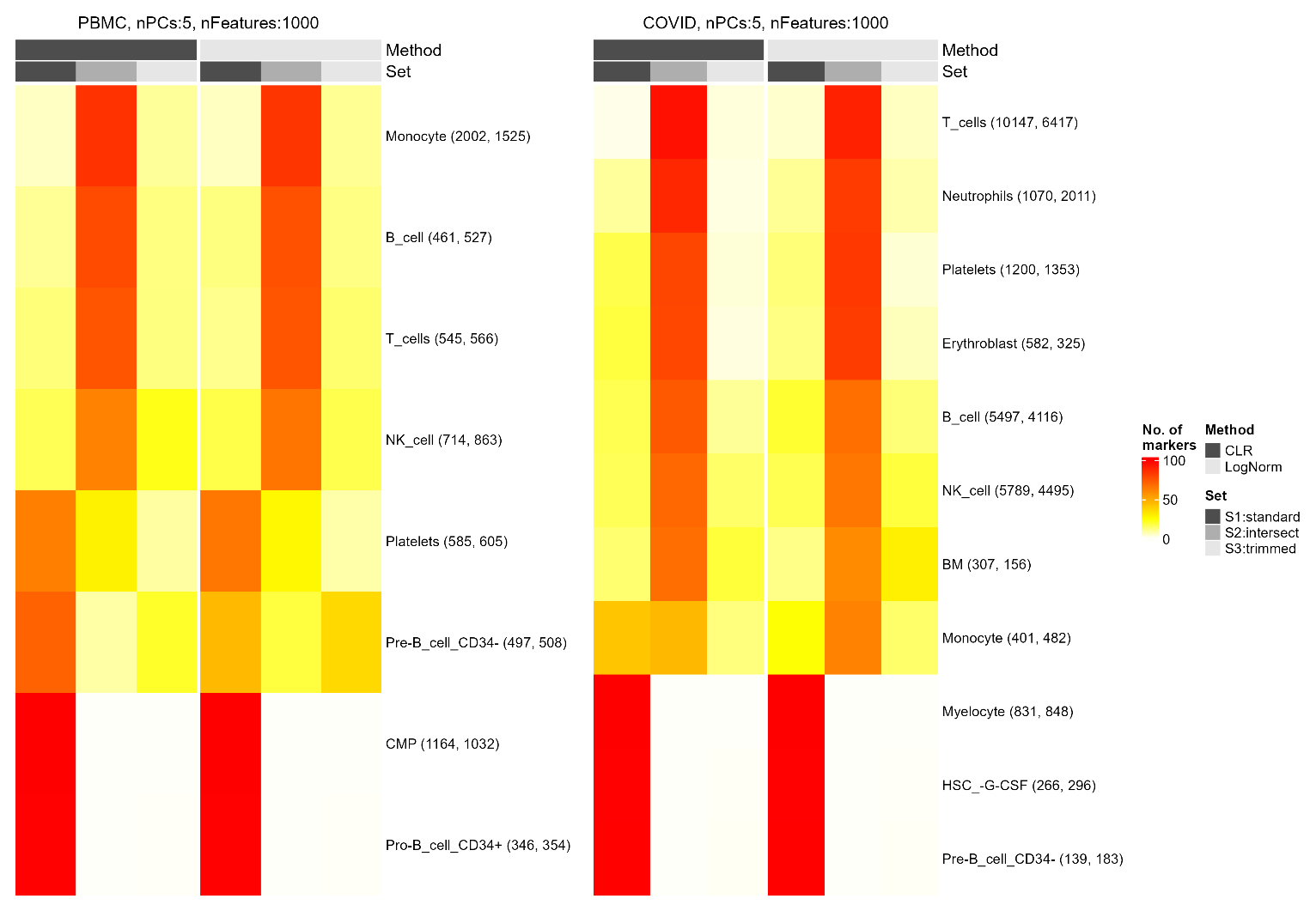
 **Supplementary Figure 2:** Heatmap comparing results for the PBMC (left) and COVID-19 (right) datasets using CLR (first three columns) or LogNorm (last three columns) normalization, where columns 1 and 4 show untrimmed-unique markers (S1: standard), columns 3 and 6 display trimmed-exclusive markers (S3: trimmed), and columns 2 and 5 represent shared markers (S2: intersect).

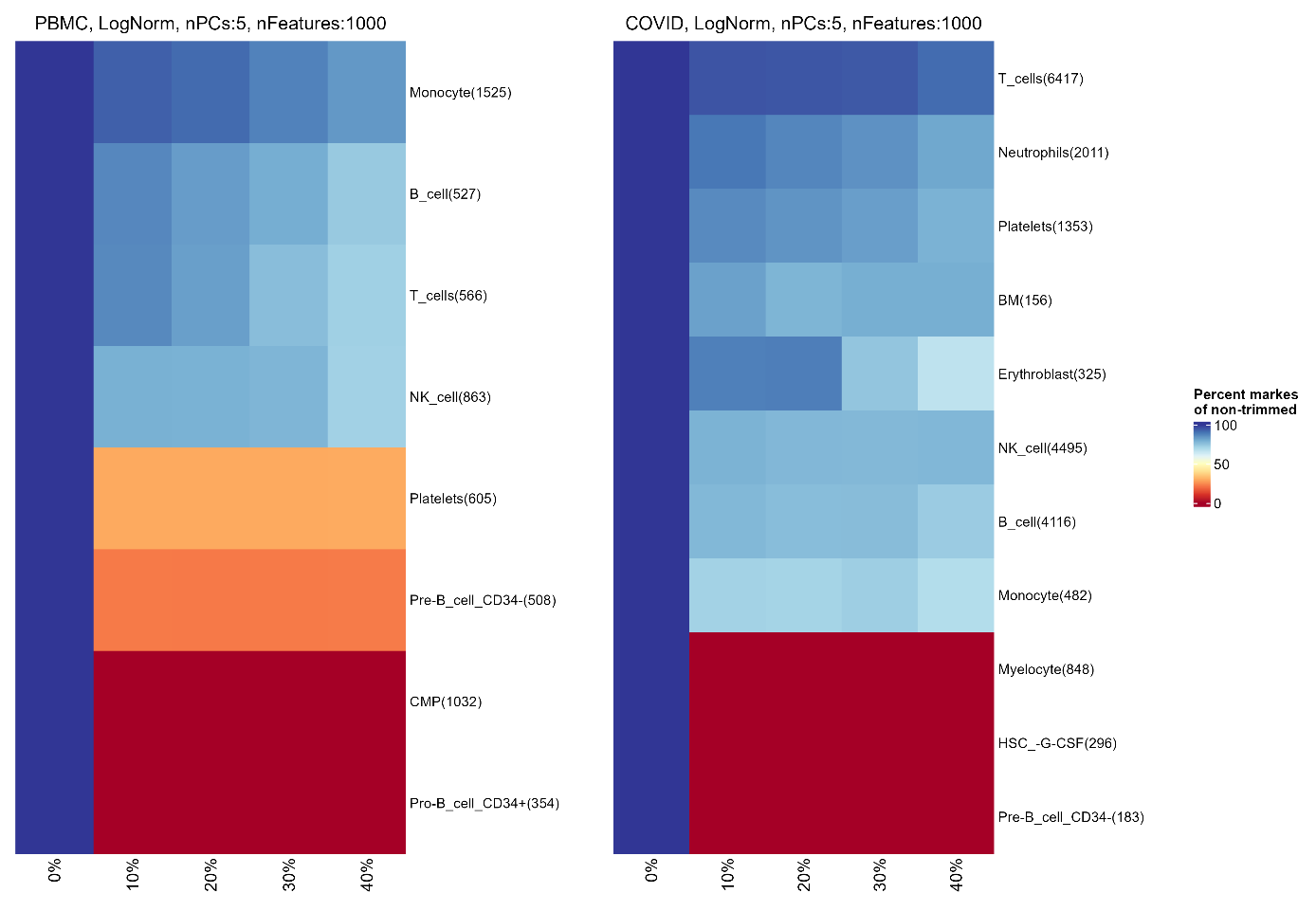

**Supplementary Figure 3:** Left: Breakdown point heatmap of PBMCs (log-normalized, 5 principal components, top 100 features). Right: COVID-19 dataset analysed under identical parameters, showing the percentage of retained marker genes (gradient from red, 0%, to blue, 100%) after trimming at 0%, 10%, 20%, 30%, and 40% thresholds, with cell type clusters labelled on the y-axis.

B) **R-Vignette for scTrimClust**

Preprocessing of input data

The identification of outlier cells for each cell type based on cluster hulls requires a preprocessed Seurat object containing a low-dimensional embedding (e.g., t-SNE or UMAP). We demonstrate this using the 2,700 Peripheral Blood Mononuclear Cell (PBMC) data set from the Seurat guided clustering tutorial ( <https://satijalab.org/seurat/articles/pbmc3k_tutorial.html> ), publicly available from 10x Genomics ( <https://cf.10xgenomics.com/samples/cell/pbmc3k/pbmc3k_filtered_gene_bc_matrices.tar.gz> ).

In short, the following preprocessing steps must be completed before using scTrimClust (here demonstrating with t-SNE as the dimensionality reduction method):

```
library(Seurat)
library(magrittr)

pbmc_sc <- Read10X("data_original/blooddata_hg19/") %>%
  CreateSeuratObject(project = "pbmc3k", min.cells = 3, min.features = 200) %>%
  {PercentageFeatureSet(., pattern = "^MT-") -> .[["percent.mt"]]; .} %>%
  subset(.,subset = nFeature_RNA > 200 & nFeature_RNA < 2500 & percent.mt < 5) %>%
  NormalizeData() %>%
  FindVariableFeatures(nfeatures = 2000) %>%
  ScaleData() %>%
  RunPCA(npcs = 10) %>%
  FindNeighbors(dims = 1:10) %>%
  FindClusters(resolution = 0.5) %>%
  RunTSNE(dims = 1:10)
```

The process includes quality control filtering, data normalization, variable feature selection, principal component analysis (PCA), and t-SNE dimensionality reduction. The preprocessed pbmc_sc seurat object contains the following components:

```
pbmc_sc
An object of class Seurat
13714 features across 2638 samples within 1 assay
Active assay: RNA (13714 features, 2000 variable features)
 3 layers present: counts, data, scale.data
 2 dimensional reductions calculated: pca, tsne
```

Gene expression profiles for 13,714 genes across 2,638 cells are stored in pbmc_sc, with 2,000 variable features within the RNA assay, three data layers (counts, data, scale.data), and computed PCA and t-SNE dimensionality reductions. Cell type annotation can be performed manually or be automated using the SingleR package, which relies on reference expression data such as the Human Primary Cell Atlas (HPCA). The HPCA data set is included in the celldex R package.

```
library(SingleR)
library(celldex)

pbmcsce <- as.SingleCellExperiment(pbmc_sc)
pbmcse <- as(pbmcsce, "SummarizedExperiment")
rownames(pbmcse) <- rownames(pbmcsce)
ref <- HumanPrimaryCellAtlasData()
pbmc_sc$CellAnnotation <- SingleR(
  test = pbmcse,
  ref = ref,
  labels = ref$label.main
)$labels
```

Here, cell type annotations are stored in the 'CellAnnotation' metadata column for use within the scTrimClust function. Alternatively, the function can utilize numeric cluster IDs from the 'seurat_clusters' column (output by FindClusters) to identify cluster-specific outliers.

Identifying outlier cells with scTrimClust

We added the approach of scTrimClust in form of three new functions to our R-package RepeatedHighDim (<https://cran.r-project.org/web/packages/RepeatedHighDim/index.html>). The core function, scTrimClust, identifies and removes outlier cells within user-defined clusters, operating on a preprocessed Seurat object containing cluster annotations (e.g., the pbmc_sc data set with labels stored in the ‘CellAnnotation’ metadata field). To maintain compatibility with Seurat, we use the package’s native DimPlot function to visualize cells during outlier detection, displaying them on dimension reduction embeddings such as UMAP, t-SNE, or PCA. scTrimClust inherits all parameters from the DimPlot function and adds new ones specific to it.

```
scTCoutput <- scTrimClust(pbmc_sc,
                       reduction = 'tsne',
                       group.by = 'CellAnnotation',
                       add.alpha.hull = TRUE,
                       hull.alpha = 2,
                       remove.outliers = FALSE,
                       outlier.quantile = 0.1,
                       outlier.alpha = 0.2)
```

The hull.alpha parameter controls the concavity of cluster boundaries using a convex hull, implemented via the ahull function from the alphahull package. Higher values produce smoother, more inclusive hulls, while lower values create tighter, more irregular contours. The outlier.quantile parameter sets the percentile cutoff (0-1) for minimum cell-to-hull distances, classifying cells below this threshold as outliers. Lower values restrict detection to extreme outliers, whereas higher values identify more peripheral cells as outliers.

Additional parameters allow further customization: add.alpha.hull allows the hull around each cluster to be added or removed, remove.outliers excludes outlier cells from both the plot and the returned Seurat object, and outlier.alpha adjusts the transparency of outlier cells when remove.outliers = FALSE. Unlike DimPlot, which returns only a ggplot object, scTrimClust returns a list of objects.

```
print(names(scTCoutput))

[1] "plot"               "object"
[3] "nonoutliers_coords" "outlier_coords"
[5] "d_ahull_coords"     "ahull_list"
```

The output includes a modified plot with flagged or removed outlier cells, along with the hull coordinates required to generate cluster boundaries (d_ahull_coords), a list of ahull objects for each cluster (ahull_list), and the coordinates for both non-outlier (nonoutlier_coords) and outlier cells (outlier_coords). It also returns the Seurat object (object), with outlier cells removed if remove.outliers = TRUE. The processed Seurat object enables seamless integration of scTrimClust into existing Seurat workflows, where DimPlot would typically be used, allowing users to continue downstream analyses without outliers.

Compare the effect of trim on marker genes for different methods

The scTC_trim_effect function quantifies changes in cluster-specific marker genes after outlier removal by comparing untrimmed (default Seurat) and trimmed (scTrimClust-processed) data sets. Assume we have already identified outliers using scTrimClust (Section 4.2), generating the trimmed Seurat object scTCoutput$object. Both pbmc_sc (untrimmed) and scTCoutput$object (trimmed) contain cell type annotations in the ‘CellAnnotation’ metadata column (Section 4.1), which we set as active identities for marker detection.

```
# Set CellAnnotation as active identity for both data sets
Idents(pbmc_sc) <- "CellAnnotation"         # Untrimmed
Idents(scTCoutput$object) <- "CellAnnotation"  # Trimmed
```

Marker gene lists from both untrimmed and trimmed data sets are required. These are generated using the FindAllMarkers function from Seurat.

markers_untrimmed **<-** FindAllMarkers**(**pbmc_sc**)**

markers_trimmed **<-** FindAllMarkers**(**scTCoutput**$**object**)**

Marker genes must be paired as lists comparing untrimmed and trimmed results. Colors for methods and gene set categories (original, shared, trimmed-exclusive) are user-defined.

```
method_pairs <- list(
  Default = list(data1 = markers_untrimmed, data2 = markers_trimmed)
)
method_colors <- c(Default = "#2CA02C")  # method color (green)
set_colors <- c(
  "S1:standard" = "#4D4D4D",   # Untrimmed-exclusive markers
  "S2:intersect" = "#AEAEAE",  # Shared markers
  "S3:trimmed" = "#E6E6E6"     # Trimmed-exclusive markers
)
```

The output is a heatmap displaying cell clusters (rows) and three categories per method (columns): markers exclusive to untrimmed data, shared markers, and markers exclusive to trimmed data. Colors represent the percentage of markers in each category (0-100%).

```
scTC_trim_effect(
  method_pairs = method_pairs,
  method_colors = method_colors,
  set_colors = set_colors,
  column_title = "PBMC Data set: Marker Changes After Trimming"
)
```

Trimming and breakdown point analysis

While scTC_trim_effect compares marker sets *across methods* (e.g., CLR vs. LogNorm) at a fixed trimming level, scTC_bpplot evaluates marker retention across trimming percentages for a single method. Trimming percentages are specified via the outlier.quantile parameter in scTrimClust. Below, we analyze the PBMC data set trimmed at 10%, 20%, 30%, and 40% (using the same normalization method, e.g., LogNorm,nfeatures = 2000, nPCs = 10), comparing marker retention to the original untrimmed (0%) data set.

We assume that the steps described in section 4.2 were already performed for 0%, 10%, 20%, 30%, and 40% trimmed. We then obtain the marker genes for each trimming level via FindAllMarkers.

pbmc_0trim **<-** FindAllMarkers**(**pbmc_sc**)**

pbmc_10trim **<-** FindAllMarkers**(**pbmc_10trimmed_object**)**

pbmc_20trim **<-** FindAllMarkers**(**pbmc_20trimmed_object**)**

pbmc_30trim **<-** FindAllMarkers**(**pbmc_30trimmed_object**)**

pbmc_40trim **<-** FindAllMarkers**(**pbmc_40trimmed_object**)**

We then use all marker gene lists as input for scTC_bpplot.

```
scTC_bpplot(
  pbmc_0trim, pbmc_10trim, pbmc_20trim, pbmc_30trim, pbmc_40trim,
  trim_percent_vector = c(0, 10, 20, 30, 40),
  legend_title = "Log-Normalized PBMC: Marker Retention Across Trimming Levels")
```

The heatmap displays cell clusters (rows, labeled with original marker counts) and trimming percentages (columns, e.g., 0%, 10%, 20%,30% and 40%), using a color gradient (default: 0% = blue, 100% = red) to show the percentage of original markers retained. High retention (warmer colors) indicates robust clusters, while cooler colors reflect marker loss after trimming.
